# A chronic interorgan wound response appropriated by *Drosophila* tumors to induce intestinal inflammation

**DOI:** 10.64898/2026.07.03.735641

**Authors:** Katy L. Ong, Jan B. Cajulao, Karina M. Anders, David Bilder

**Affiliations:** Department of Molecular and Cell Biology, University of California-Berkeley, Berkeley CA, 94720, USA

## Abstract

Tumors exploit wound healing pathways not only to foster progression but also to lethally disrupt systemic physiology. A prominent example is malignant activation of the clotting cascade, causing pathology through unclear mechanisms that extend beyond thrombosis. Here we show that tumors in a coagulopathy-inducing *Drosophila* cancer model remotely disrupt intestinal stem cell (ISC) homeostasis. This paraneoplastic, tumor-gut communication axis induces intestinal dysplasia and barrier dysfunction, mimicking remote chronic injury that we show activates inflammation in ISCs via EGFR signaling. Unlike the interorgan responses observed with acute injury, which involves Jak/STAT signaling to activate regenerative ISC proliferation, dysregulated division in both tumor-bearing and chronically injured flies is sustained only by EGFR activation. We also present evidence that deposition of clot material locally onto gut stroma links tumor-driven coagulopathy to intestinal inflammation. Collectively, these findings distinguish mechanisms of remote responses to chronic versus transient stresses. Furthermore, they show how tumor-initiated, dysregulated wound healing programs can drive tissue-specific, pathological inflammation in a host.

## INTRODUCTION

As the ‘emperor of maladies’, cancer introduces chaos into the body at many biological levels. Much of the effort to mitigate cancer’s harm has focused on the tumor mass effect, invasion and metastasis, all of which disrupt tissues through direct contact (Hanahan, 2022; Hanahan & Weinberg, 2011).

However, pathological interactions can occur in the absence of a physical interface, resulting in systemic dysregulations of physiology that are sometimes grouped as paraneoplastic syndromes (McAllister & Weinberg, 2014; Sardiña González *et al*, 2022). Such systemic effects remain under-addressed in cancer biology despite accounting for a substantial portion of its disease burden. For example, cachexia is estimated to directly cause up to 20% of cancer patient deaths, but its underlying mechanisms are only now yielding to experimental investigation (Tisdale, 2002, 2009; Fearon *et al*, 2013; Bruera, 1997). In the past decade, *Drosophila* tumor models have contributed to untangling complex but conserved tumor-host communication axes, including those that cause cachexia (Kwon *et al*, 2015; Figueroa-Clarevega & Bilder, 2015; Saavedra & Perrimon, 2019; Song *et al*, 2019; Ding *et al*, 2021; Lodge *et al*, 2021; Bilder *et al*, 2021). The experimentally tractable, *in vivo* fly platform has further identified candidate pathways and provided insights into mechanistic bases of other human paraneoplasias (Yeom *et al*, 2021; Cong *et al*, 2025; Kwok *et al*, 2024; Xu *et al*, 2024, 2023; Hsi *et al*, 2023).

Among the physiological systems impacted by cancer, a well-established instance is systemic interference with the clotting cascade, causing lethal coagulopathies that present a major clinical challenge. Pathological thrombosis is estimated to kill up to 9% of cancer patients directly, and hijacking of the clotting cascade has been implicated in tumorigenesis and metastasis (Donnellan & Khorana, 2017; Khorana *et al*, 2007; Boccaccio & Comoglio, 2009). Yet beyond these known thrombotic mechanisms, dysregulated clotting remains correlated with poor prognosis. Patients with subclinical elevation in clotting markers have reduced survival rates, revealing a gap in knowledge of how coagulopathies kill patients in addition to thrombosis and tumor progression (Marchetti *et al*, 2014; Krenn-Pilko *et al*, 2015; Pichler *et al*, 2013; Green *et al*, 1992).

Harold Dvorak’s seminal review conceptualizing tumors as “wounds that do not heal” articulated how an understanding of the wound response could inform oncology (Dvorak, 1986). Dvorak first drew parallels between wounds and tumors in the remodeling of stromal tissue, but this perspective has since been extended to cell proliferation, neovascularization and immune interactions (Dvorak, 2015; Schäfer & Werner, 2008; Balkwill & Mantovani, 2001; Coussens & Werb, 2002). Logic follows that we can gain insight into paraneoplastic phenomena by comparative study with systemic responses to wounds. In contrast to the adaptive process of wound healing, the responses provoked by tumors are pathological. How these adaptive and pathological mechanisms are molecularly and cellularly distinguishable remains a critical open question.

We previously developed an ovarian tumor model in *Drosophila* and showed that it exhibits blood clotting phenotypes that mirror cancer-associated coagulopathies, including enhancement of host mortality (Hsi *et al*, 2023). Because flies have an open circulatory system and are not vulnerable to thrombosis, this model allows investigation of new mechanisms by which clotting dysregulation disrupts host physiology. Here, we report that tumor-driven coagulopathy aggravates an inflammatory interorgan communication axis. In the fly model, we reveal that tumors stimulate intestinal stem cell (ISC) division to the point of causing gut dysplasia and barrier failure. EGFR signaling is required for tumor-driven activation of ISCs, in a coagulopathy-dependent fashion. ISC proliferation in tumor-bearing animals is independent of Jak/STAT signaling, in contrast to proliferation triggered by acute epithelial injury. Instead, this tumor-driven process parallels a remote, chronic wound response that is observed in a new persistent injury model. Overall, this study uncovers a systemic signaling mechanism underlying pathological responses to tumors that shares features with chronic injury response, differentiating it from those that respond adaptively to acute injury.

## RESULTS

### Tumors remotely trigger intestinal dysplasia and gut barrier failure

We previously reported that genetically induced ovarian tumors cause *Drosophila* hosts to die ∼50% earlier than control counterparts (**Fig. 1A**) (Hsi *et al*, 2023). Inflammatory opening of the blood-brain barrier accounts for part but not all of this lethality, suggesting the presence of additional, unidentified lethal mechanisms(Kim *et al*, 2021). Studies in *Drosophila* identify dysfunction of the gut barrier as an early feature of impending death from multiple causes; this principle may extend to vertebrates (Cansell *et al*, 2025; Salazar *et al*, 2023; Dambroise *et al*, 2016). To examine whether intestinal failure could contribute to early lethality in the ovarian carcinoma (OC) model, we examined the cellular makeup of guts in tumor-bearing flies. Compared to control animals, the proportion of ISCs and progenitor enteroblasts (EBs) that express the marker *escargot* (*esg^+^*) substantially increased.

**Figure 1:**
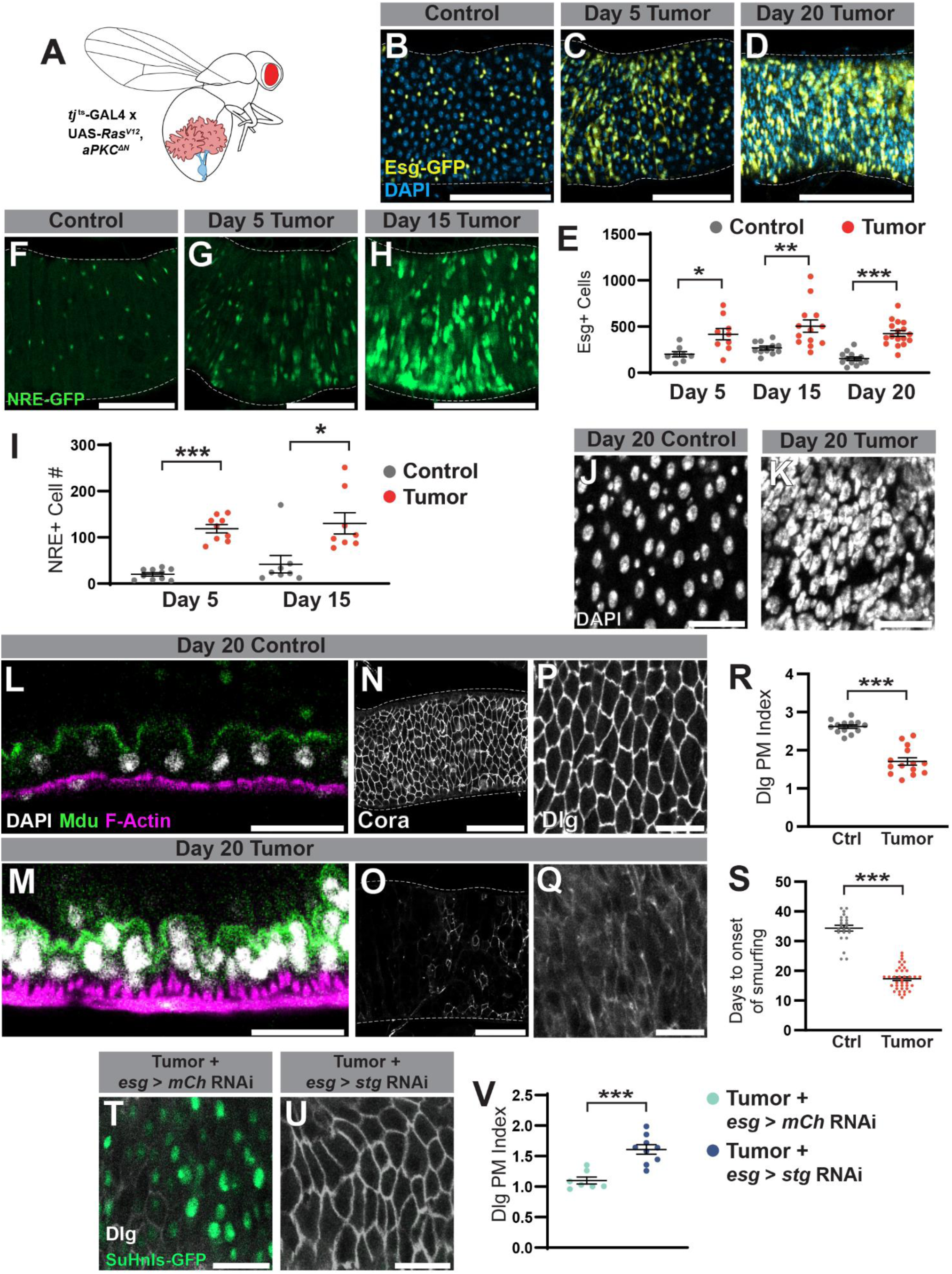
Tumors dysregulate intestinal stem cell homeostasis from a distance. (**A**) Schematic of genetic OC model in *Drosophila*. (**B**-**E**) Intestines of tumor-bearing flies (**C, D**) exhibit expansion of *esg+* cells (yellow) compared to control animals (**B**), quantitated in (**E**). (**F**-**I**) EB (green) production is elevated in tumor-bearing animals (**G, H**) compared to controls (**F**), quantitated in (**I**). (**J, K**) A dramatic shift in nuclear density is observed in tumor-bearing fly intestines 20 days ATI. (**L**-**M**) Cross-section of epithelium reveals multi-layering and disorganization in tumor-bearing animals. (**N**-**R**) SJs marked with (**N**, **O**) Coracle and (**P**, **Q**) Dlg become disorganized in OC flies, Dlg plasma membrane association quantitated in (**R**). (**S**) Smurf assays shows early gut barrier failure in tumor-bearing flies. (**T**-**V**) Inhibition of ISC division in tumor-bearing animals (**U**) depletes progenitor production and partially rescues SJ organization, quantitated in (**V**). Scale bars in **B**-**D**, **F**-**H** = 100µM. Scale bars in **J**, **K**, **L**-**Q**, **T**, **U**= 25µM. Error bars = S.E.M. Student’s t-test used to assess statistical significance in all graphs.

This started at day 5 after tumor induction (ATI) and continued to day 20, which is the approximate median time of death for tumor-bearing animals (**Fig. 1B-E)** (Hsi *et al*, 2023). Differentiated enterocytes (*esg-,* large-nucleated cells), the predominant gut cell type under homeostasis, were outnumbered by the expansion of *esg+* cells by day 20 ATI (**Fig. 1D**). To determine if increased progenitor production could account for the change in cellular composition, we analyzed a Notch activity reporter that is specifically expressed in EBs but not ISCs. This revealed a dramatic increase in the EB population of OC flies (**Fig. 1F-I)**, resulting in elevation of cellular density (**Fig. 1J, K**). In addition to cellular makeup, the guts of tumor-bearing flies exhibited marked changes in morphology. Cross-sections of the epithelium revealed loss of monolayer organization of cells and increased thickness of tissue (**Fig. 1L, M**). All of these phenotypes are reminiscent of the hyperproliferative dysplasia observed in aging as well as conditions that cause local intestinal injury and inflammation (Biteau *et al*, 2008; Jiang *et al*, 2009; Amcheslavsky *et al*, 2009; Cronin *et al*, 2009; Buchon *et al*, 2009b).

We asked how the alterations in the gut epithelium affect its function in segregating the intestinal lumen from the rest of the body. In tumor-bearing flies, a severe disorganization of septate junctions (SJs, the insect equivalent of tight junctions) was observed. The SJ markers Discs Large (Dlg) and Coracle were no longer concentrated along the plasma membrane but instead diffuse across the cytoplasm (**Fig. 1N-R**). This phenotype has been previously documented in flies that experience breaches in their intestinal barrier, shown by increased intestinal permeability using the Smurf assay (Resnik-Docampo *et al*, 2017; Clark *et al*, 2015). Consistent with this, we found an earlier onset of smurfing in tumor-bearing hosts compared to controls (**Fig. 1S**).

To test whether barrier integrity loss is caused by hyperproliferation within the gut, we inhibited ISC division in tumor-bearing flies. To accomplish this, we switched to an allograft model where host cells can be genetically manipulated independently of ovarian oncogene expression. Ovarian tumors were harvested from donors and individual transformed ovarioles were transplanted into host flies where they grafted and grew (**Fig. S1A, B**). Host flies develop paraneoplastic phenotypes comparable to OC flies, including cachectic wasting of the fat body, bloating, loss of late-stage clotting function (i.e. melanization), and a shortened lifespan (**Fig. S1C-I**). Importantly, they also exhibit dysregulation of ISC homeostasis and SJ disorganization in the intestinal epithelium (**Fig. S1J-M**).

To inhibit ISC division, RNAi against the *cdc25* homolog *string* (*stg*) was driven with temperature controlled *esg-GAL4* in adult hosts that received ovarian tumor transplants. As expected, this manipulation acutely depleted the number of EBs as compared to tumor-bearing flies expressing a control RNAi (**Fig. 1T, U**). Importantly, it also substantially rescued epithelial SJ organization (**Fig.1T-V)**. Thus, overactivation of stem cells is an underlying cause of paraneoplastic gut barrier failure.

### Cachexia, anorexia and microbiome changes do not account for disrupted ISC homeostasis

As in cancer patients, tumors in *Drosophila* can induce paraneoplastic dietary and metabolic pathologies. We investigated whether these effects might account for the gut phenotypes observed in OC flies. First, we inhibited cachexia by depleting the cachetogenic insulin antagonist ImpL2 in tumor tissue (Kwon *et al*, 2015; Figueroa-Clarevega & Bilder, 2015). We have previously reported that *ImpL2* knockdown in OC tumors did not improve host survival (Hsi *et al*, 2023; Kim *et al*, 2021).

Consistent with this, no rescue of SJ integrity or cell density was detected with this manipulation (**Fig. S2A-C**). Overexpression of ImpL2 in the ovarian follicle without tumor induction was also not sufficient to increase the number of gut progenitors labelled with STAT-GFP (**Fig. S2D, E**). Second, because starvation inhibits ISC division, we considered whether anorexia could impact digestion in tumor-bearing flies (O’Brien *et al*, 2011). Both CApillary FEeding (CAFE) and Excreta Quantification (EX-Q) assays indicated a small decrease in consumption in OC compared to control flies at 5 days ATI but no significant differences at subsequent time points (**Fig. S2F, G**). Lastly, since both enteric infection and age-related dysbiosis can cause inflammatory dysplasia, intestinal phenotypes could be a consequence of alterations in the gut microbiome (Guo *et al*, 2014; Cronin *et al*, 2009; Jiang *et al*, 2009; Buchon *et al*, 2009b; Clark *et al*, 2015). This has been observed in an allograft tumor model in the adult fly (Cong *et al*, 2025). However, we did not find a significant change in bacterial loads in OC flies compared to age-matched controls (**Fig. S2H**). Moreover, treating OC flies with antibiotic cocktails to eliminate the gut microbiome did not reinstate SJ integrity nor ISC homeostasis (**Fig. S2I-L**). Collectively, the data fail to support microbial dysbiosis, anorexia or cachexia as drivers of dysplasia and gut barrier failure in OC flies. They instead point towards an alternative mechanism for increased ISC activity in the tumor-bearing flies.

### Chronic injuries also stimulate a remote wound response in the intestine

We considered whether an ongoing inflammatory process, perhaps related to that induced by tissue damage, could trigger dysplasia. Local insults to the gut such as ingested pathogenic bacteria or toxins are well-known to stimulate ISC division (Cronin *et al*, 2009; Jiang *et al*, 2009; Buchon *et al*, 2009b; Amcheslavsky *et al*, 2009). The gut can also respond to distant wounds at the fly’s surface: following a clean injury to the thorax or abdomen, ISC proliferation is stimulated (Chakrabarti *et al*, 2016; Chakrabarti & Visweswariah, 2020; Takeishi *et al*, 2013). In these responses to transient damage, the gut maintains and/or restores homeostatic character, preserving the normal proportion of EBs (Chakrabarti *et al*, 2016; Takeishi *et al*, 2013; Buchon *et al*, 2010; Jiang *et al*, 2011). We wondered whether a wound response that was pathologically sustained might have a different outcome, closer to that resulting from a tumor. To induce persistent damage in a distant tissue, we used two genetic strategies. First, the pro-apoptotic gene *reaper (rpr)* was conditionally overexpressed to induce death in follicle epithelial cells (**Fig. 2A, Fig. S3A, B**). Second, the matrix metalloprotease MMP1 was transiently induced in follicles to disrupt the basement membrane.

**Figure 2.**
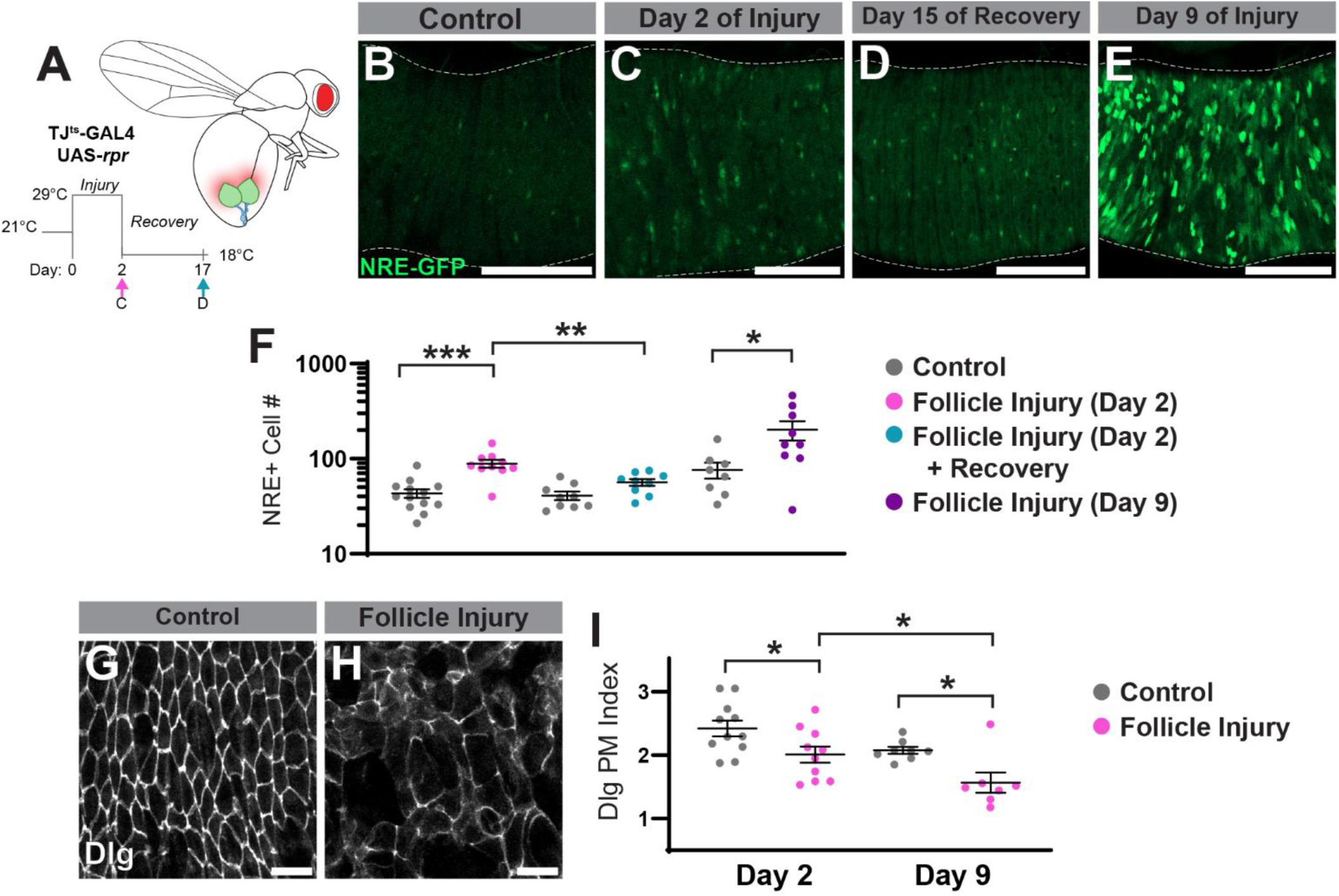
Chronic follicle injury triggers a tumor-like, systemic response in the gut. (**A**) Schematic of ovarian injury timeline and experimental procedure. (**B**, **C**) NRE-GFP reporter revealed EB elevation following two days of ovarian injury. (**D**) Repression of apoptosis following injury attenuates EB numbers. (**E**) Chronic injury for 9 days leads to a larger increase in EB population. Quantification in (**F**). (**G**-**I**) Compared to controls (**G**), chronic ovarian injury leads to disruption of SJ organization (**H**), quantified in (**I**). Scale bars in **B**-**E** = 100µm. Scale bars in **G**, **H** = 25µm. Error bars = S.E.M. Student’s t-test used to assess statistical significance in all graphs.

Remarkably, we found increased EB counts in both genotypes, demonstrating that, in contrast to acute injuries, chronic-type injuries in a distal tissue can disrupt gut stem cell homeostasis (**Fig. 2B, C, F, Fig. S3C-E**).

We then wondered if the chronic wounding response would abate as distant damage was resolved. When *rpr* expression for 2 days was subsequently restricted to allow the ovary to recover, EB numbers declined albeit to a level slightly higher than that of control flies, indicating that the gut reaction is semi-reversible (**Fig. 2A, D, F**). By contrast, when the injury was allowed to persist for 9 days of *rpr* expression, the EB population expanded to numbers resembling those seen in OC flies (**Fig. 2E, F**). Moreover, persistent follicle epithelial death resulted in a decrease in intestinal SJ organization, evident two days after ovarian injury and becoming more severe 9 days post-injury (**Fig. 2G-I**). These results align with the hypothesis that OC tumors act like unhealable wounds and provoke a chronic injury response that includes pathological communication to the gut.

### EGFR overrides other stress signaling pathways to drive intestinal dysplasia in tumor-bearing and chronically injured flies

The data above suggest that tumors overstimulate a normally adaptive, inflammatory response. We thus asked which injury-activated inflammatory pathways might be responsible for loss of ISC homeostasis. We first examined the Jak/STAT pathway. IL-6-like cytokine signaling drives age-related intestinal dysplasia as well as responses to both distant and local acute injury (Guo *et al*, 2014; Cronin *et al*, 2009; Jiang *et al*, 2009; Chakrabarti *et al*, 2016; Buchon *et al*, 2009a; Li *et al*, 2016). Additionally, OC tumors upregulate several IL-6-like Unpaired proteins, which activate inflammatory signaling that compromises the blood-brain barrier (Hsi *et al*, 2023; Kim *et al*, 2021). As expected, ovarian tumors resulted in elevated STAT activity in the intestinal epithelium, and we further found that this was true with chronic ovarian injury (**Fig. 3A-C, Fig. S3F**-**H**). To test whether STAT activation was required for paraneoplastic ISC hyperproliferation, we transplanted ovarian tumors into flies in which Stat92E or the STAT pathway receptor Domeless was depleted in *esg^+^* cells.

**Figure 3.**
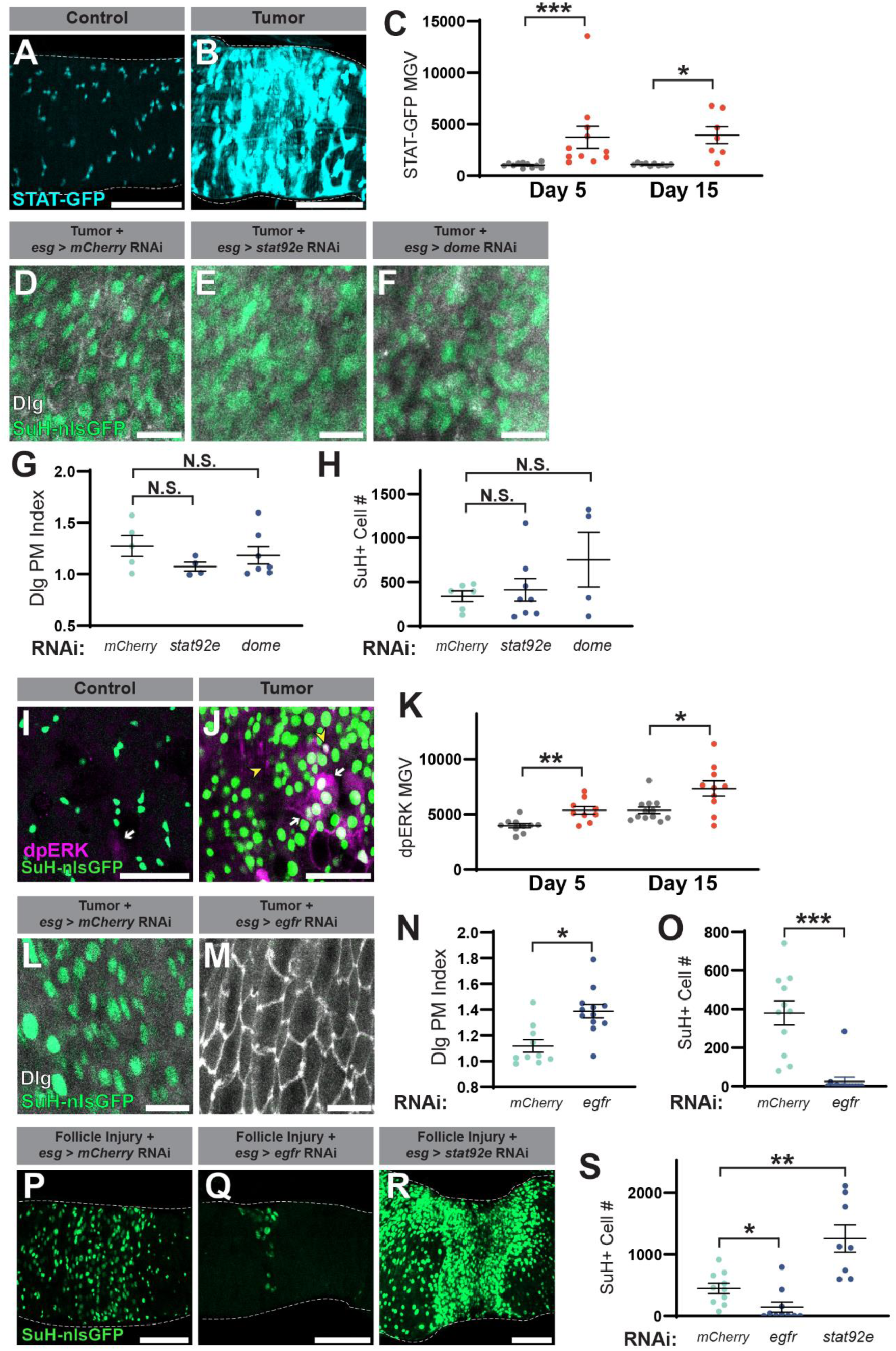
EGFR overactivation drives tumor-driven gut dysplasia, bypassing Jak/STAT control of ISCs. (**A**-**C**) STAT-GFP reporter (cyan) reveals pathway activation in the gut upon ovarian tumor induction. Quantification in (**C**). (**D**-**H**) Knockdown of JAK/STAT pathway components in the ISCs/EBs did not rescue SJ integrity (white) or EB expansion (green). Quantification of Dlg plasma membrane association in (**G**), EB cell number in (**H**). (**I**-**K**) dpERK immunofluorescence (magenta) reveals MAP Kinase activation in small– (yellow arrowheads) and large-nucleated (white arrows) intestinal cells with ovarian tumors. Quantification in (**K**). (**L**-**O**) *EGFR* knockdown in ISCs/progenitors rescues SJ integrity and EB expansion. Quantification of Dlg plasma membrane association in (**N**), EB cell number in (**O**). (**P**-**S**) *EGFR* but not *Stat92E* knockdown rescues excess EBs generated upon chronic follicle injury. Quantification in (**S**). Scale bars in **A**, **B**, **P**-**R** = 100µm. Scale bars in **D**-**F** = 20µm. **I**, **J** = 50µm. **L**, **M** = 25µm. Error bars = S.E.M. Student’s t-test used to assess statistical significance in all graphs.

Surprisingly, neither manipulation mitigated the excess number of EBs in the epithelium nor restored the loss of SJ integrity (**Fig. 3D-H**). We also combined dual expression systems in order to determine the requirement for Jak/STAT in the gut during chronic injury of the follicle epithelium. *LexAop-Rpr* was expressed under temperature controlled *tj-LexA::GAD* to kill follicle cells while Stat92E was simultaneously depleted by RNAi with *esg-Gal4*. Not only was no decrease in EBs observed, but this immature cell population was slightly elevated, possibly due to defective differentiation with *stat92e* knockdown (Beebe *et al*, 2010) (**Fig. 3P, R, S**). Thus, in contrast to the response to acute injury, inflammatory cytokine signaling is not essential for tumor-induced ISC activation.

Since cytokine signaling was not required, we sought alternative pathways that could alter ISC proliferation. EGFR signaling functions in parallel with Jak/STAT to control stress-induced ISC division (Buchon *et al*, 2010; Jiang *et al*, 2011). In response to damage, several midgut cell types upregulate EGF ligands and signaling proteases to initiate compensatory proliferation (Buchon *et al*, 2010; Jiang *et al*, 2011). To probe whether this pathway in the intestinal epithelium might be remotely stimulated by tumors or chronic wounds, we analyzed levels of phosphorylated ERK (dpERK) which is generated by MAP Kinase following EGFR activation. We found that, for both conditions, elevated MAP Kinase activity was observed in small-nucleated cells (ISCs, progenitors and/or neuroendocrine cells), large-nucleated enterocytes, and in the visceral muscle layer (**Fig. 3I-K, Fig. S3I-K).** To test whether EGFR signaling in ISCs generates dysplasia in response to ovarian tumors, we depleted EGFR in tumor-bearing hosts. Upon knockdown of EGFR in ISCs and progenitors, both ISC proliferation and SJ disorganization in response to tumor grafting were substantially mitigated (**Fig. 3L-O**). Additionally, *egfr* knockdown with chronic follicle injury substantially reduced the EB population (**Fig. 3P, Q, S**). These data identify a central role for EGF signaling in both tumor– and chronic-injury induced ISC proliferation; activation of this pathway supersedes other proliferation control mechanisms in the epithelium such as Jak/STAT.

### Tumor-driven coagulopathy promotes a chronic inflammatory state selectively in the gut

Paraneoplastic effects can be regulated directly by tumor-secreted oncokines, and OC tumors are known to produce several such factors. However, of the four *Drosophila* EGF ligands, only Vein is transcriptionally upregulated in the transformed ovary (Hsi *et al*, 2023). Because the Neuregulin-like Vein is the least potent of the four and is generally used for short-range signaling (Schnepp *et al*, 1996; Golembo *et al*, 1999; Schnepp *et al*, 1998), we investigated other oncokines that might activate gut EGFR signaling indirectly.

Our prior work unveiled a role for tumor-secreted clotting proteins in coagulation defects and impaired wound healing, as well as early mortality of the host. However, the connection between clotting dysregulation and tumor-induced physiology changes was not determined. We examined gut dysfunction when various clotting components were depleted in the tumor. RNAi-mediated knockdown of each of three different clotting factors –– Fon, Fbp1 and Eig71ee –individually rescue wound healing defects (**Fig. 4SA, B**) (Hsi *et al*, 2023; Bajzek *et al*, 2012; Lindgren *et al*, 2008; Balasubramanian *et al*, 1998). Strikingly, each also led to a significant attenuation of SJ integrity accompanied by an extension of host lifespan (**Fig. 4A-D, Fig. S4C**) (Hsi *et al*, 2023). Rescue of these phenotypes occurs without alteration of tumor progression (**Fig. S4D, E**), consistent with a paraneoplastic mode of action (Hsi *et al*, 2023). *fon* depletion in the tumor with two different RNAis significantly delayed the onset of smurfing (**Fig. 4E, Fig. S4F**); for the remainder of this study, we used *fon* depletion to inhibit tumor-driven coagulopathy. These data suggest that tumor-driven coagulopathy participates in interorgan signaling to cause lethal gut dysfunction.

**Figure 4.**
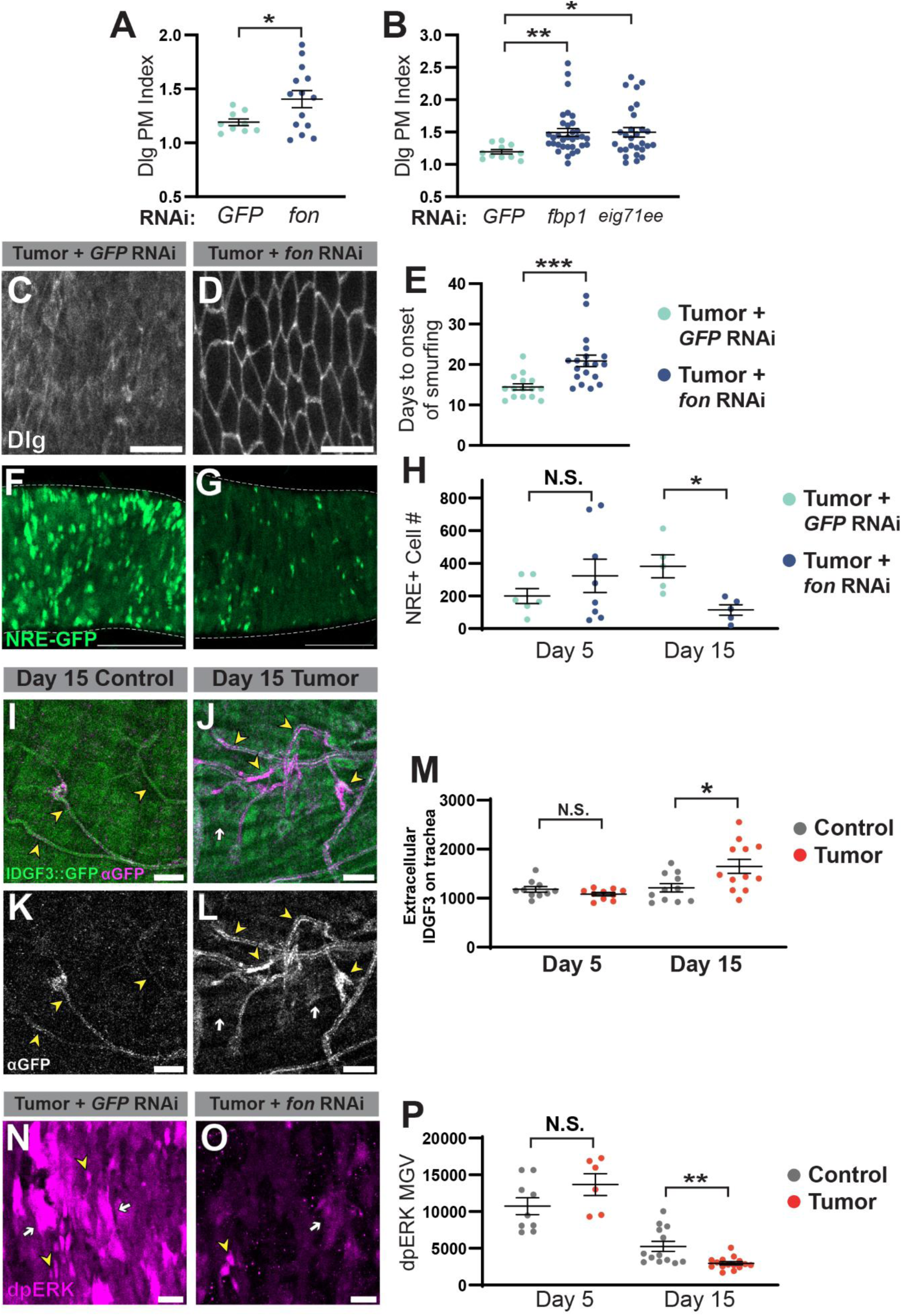
Tumor-driven coagulopathy is required for chronic ISC activation. (**A**-**D**) Knockdown of clotting components in the tumor rescues SJ integrity (white) in the gut. (**E**) Knockdown of *fon* in the tumor delays the onset of smurfing. (**F**-**H**) Rescue of EB expansion (green) is observed at day 15 ATI with *fon* knockdown in tumor cells. Quantification in (**H**). (**I**-**M**) The clotting component IDGF3 is recruited to the visceral muscle (white arrows) and gut-associated trachea (yellow arrowheads) in the context of an ovarian tumor. Green shows intrinsic IDGF3::GFP signal. Magenta and white display extracellular pool of anti-GFP staining. Quantification of trachea-associated signal in (**M**). (**N**-**P**) dpERK immunofluorescence (magenta) reveals attenuation of MAP Kinase activity in small– (yellow arrowheads) and large-nucleated (white arrows) upon *fon* knockdown in tumor cells 15 days ATI, quantification in (**P**). Scale bars in **C**, **D** = 25µm. Scale bars in **F**, **G** = 100µm. Scale bars in **I**-**L**, **N, O** = 20µm. Error bars = S.E.M. Student’s t-test used to assess statistical significance in all graphs.

We explored the kinetics of this involvement. Early stages of tumor-induced gut dysplasia (day 5 ATI) showed no difference in EB numbers between *fon* RNAi and control conditions (**Fig. 4H**). However, a substantial reduction in EBs was seen with *fon* knockdown at later stages (day 15 ATI) (**Fig. 4F-H**).

This experiment indicates different phases of ISC activation governed by separable processes, and that coagulopathy interfaces with the later phase of gut inflammation.

**Figure 5.**
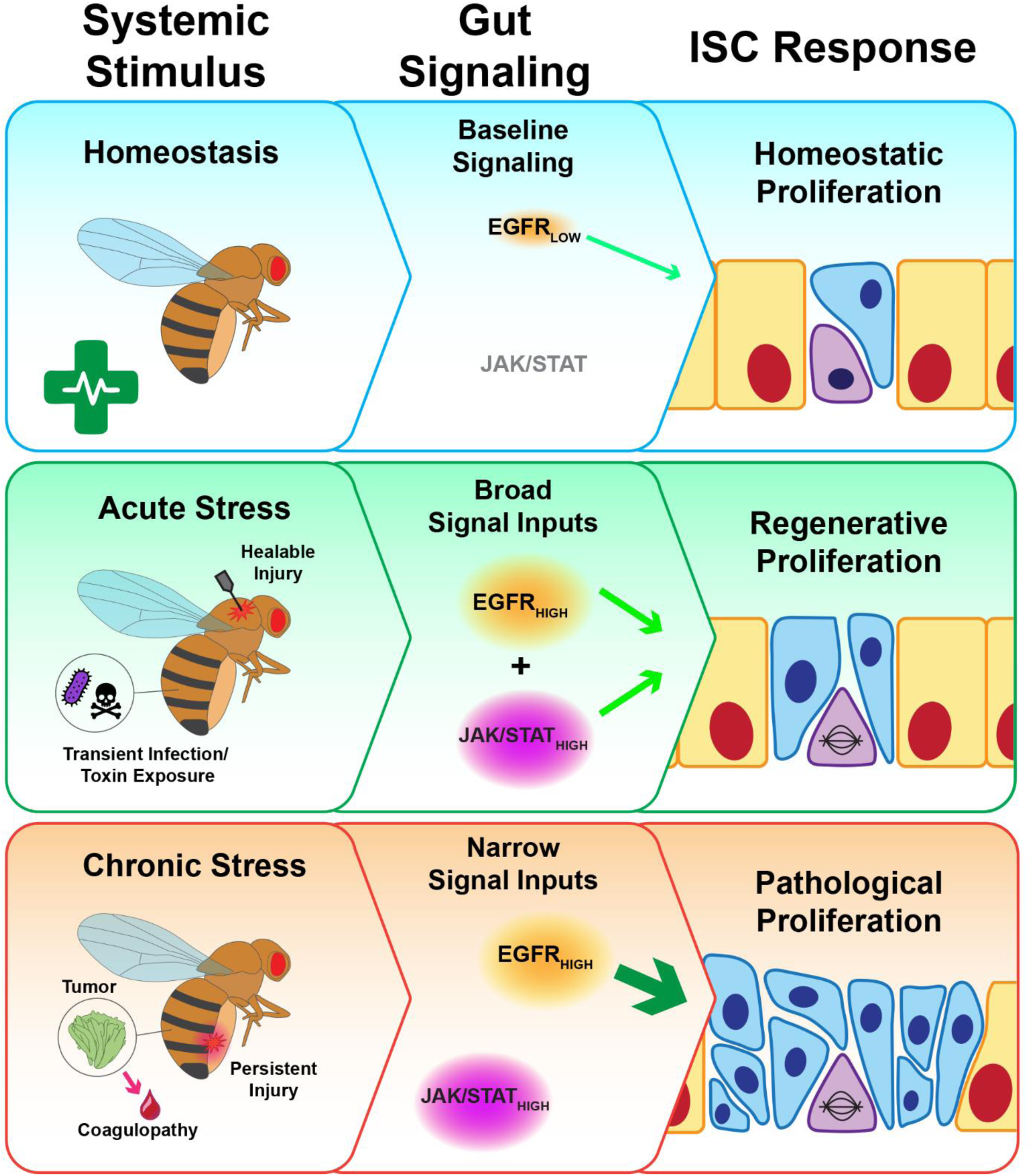
Malignancy and chronic injury share a distinct signaling axis to induce remote inflammation.

To explore this interface, we assessed the localization of a clotting component (IDGF3) that is endogenously tagged with GFP (Kucerova *et al*, 2015). Using antibody staining without detergent to visualize extracellular species, IDGF3-GFP aggregates were notably detected on gut-associated trachea (visible in the GFP channel due to auto-fluorescence) in tumor-bearing as compared to control flies, and to a lesser extent on visceral muscle (**Fig. 4I-M**). Elevation of tracheal signal was observed at day 15 but not day 5 ATI, correlating with the time point when *fon* knockdown in the tumor rescues gut phenotypes (**Fig. 4M**). Extracellular aggregates on trachea were also seen when transgenic Fon::GFP was overexpressed in ovarian tumors (**Fig. S4G-J**). These signals were difficult to quantify because of their overlap with the strong muscle-associated signals previously documented(Green *et al*, 2016), but qualitatively, tracheal Fon::GFP appeared enhanced in tumor-bearing flies (**Fig. S4I, J**). Overall, the accumulation of clotting components directly on gut stromal tissues in tumor-bearing flies suggest that, as in vertebrates, the fly clotting reaction may be catalyzed on vascular structures, raising the possibility that this may mechanistically link coagulopathy to chronic activation of ISCs.

Finally, given the critical roles for both EGFR signaling and clotting in tumor-driven gut dysplasia, we assessed whether MAP Kinase activation was affected by coagulopathy. Indeed, *fon* knockdown in the tumor substantially decreased dpERK levels in the intestinal epithelium compared to tumor-specific knockdown of a control RNAi (Fig. **4N-P**). Consistent with the hypothesis above, the difference was seen at day 15 but not day 5 ATI (**Fig. 4P**). Tumor-specific *fon* knockdown also resulted in decreased STAT reporter activity in the gut but not the tumor, muscle, or dorsal tissues of OC flies **(Fig. S4K-M)**, pointing to a tissue-specific role in intestinal inflammation. Together, these data suggest a model in which tumor-driven coagulopathy, through local action on intestinal stroma, potentiates chronic EGFR signaling to increase ISC activity and intestinal barrier failure.

## DISCUSSION

The true extent of communication through the ‘internal milieu’ that maintains homeostasis in animal bodies, and how it is pathologically disrupted, is only now being elucidated (Leopold & Perrimon, 2007). Over the past 12 years, *Drosophila* tumor models have revealed normal and paraneoplastic signaling to fly analogs of human liver, adipose, kidney, brain and immune tissue (Bilder *et al*, 2021). Our data here extend these axes to the gut, which is an emergently central regulator of organism viability (Salazar *et al*, 2023). Cong et al. recently demonstrated that renal damage caused by a transplanted fly tumor is secondary to gut dysbiosis and sepsis, but did not describe a mechanism for gut leakiness (Cong *et al*, 2025). Here we show that a different fly tumor can remotely elicit gut barrier loss independent of microbiome changes through EGFR-stimulated proliferation of ISCs. The EGF ligand that triggers ISC activation is not identified in the present work. Previous studies of homeostatic conditions demonstrate that three ligands from different gut cell types act largely redundantly (Jiang *et al*, 2011; Biteau & Jasper, 2011); whether the same ligands regulate both physiological and pathological EGFR activity remains an open question.

Our data shine light on comparisons between acute injury, chronic wounds, and tumors, with a special focus on systemic responses. Reactions in the gut were similar when insult was caused by either malignant transformation or chronic damage in a remote tissue: both processes relied on strong intestinal EGF signaling to trigger dysplastic proliferation. This contrasts with acute injury to the distant surface epithelium as well as to the gut itself, where Jak/STAT was dispensable for tumor– and chronic injury-stimulated EB expansion. The results recall previous ectopic activation studies showing that EGFR can drive proliferation in the absence of JAK/STAT signaling (Buchon *et al*, 2010; Jiang *et al*, 2011). Overall, they align with the hypothesis that tumors and chronic wounds activate potent EGFR signaling to pathologically rather than homeostatically promote ISC division. Thus, the systemic reactions elicited by tumors can be distinguished from the adaptive responses to acute, resolvable injury. Following in the steps of Dvorak, comparative study of malignancy and chronic injury responses could elucidate further mechanisms of systemic pathophysiology.

A provocative facet of this work is evidence that tumor-driven coagulopathy acts as an accelerant of inflammation in the fly gut. In humans, blood clotting plays roles in both initiation and resolution of inflammation. Could hyperinflammation contribute to the mortality burden of cancer coagulopathies? Gut barrier integrity is often compromised in cancer patients (Wardill & Bowen, 2013; Klein *et al*, 2013; Yu & Schwabe, 2017). Chemotherapy and microbiome alterations may not fully account for this defect, with studies pointing to the involvement of other unknown factors (Eiman *et al*, 2025; Puppa *et al*, 2011; Klein *et al*, 2013). Tumor-triggered coagulopathy could be such a factor, and indeed dysregulated clotting is associated with multiple inflammatory bowel diseases (Danese *et al*, 2007; Lagrange *et al*, 2021). Given the absence of pathological thrombosis in flies and the role for tumor-driven coagulopathy in inflammatory gut damage and lethality, *Drosophila* is well poised to explore this hypothesis.

Our report leaves open an intriguing question of how coagulopathy is mechanistically linked to tissue-specific physiology changes. In mammals, clots incorporate secreted factors with signaling potential, including growth factors (Stachowicz *et al*, 2017; Scridon, 2022). An attractive hypothesis is thus presented in which tumor-driven coagulopathy promotes a local concentration increase of EGF ligands on fly gut stroma, enabling remote physiological changes of tissue-specific host organs by cancer cells. Such a result would further extend the significant analogies between clotting system of insects and humans despite their having evolved largely independently. Indeed, our studies highlight the surprising amount of knowledge to be gained from comparative studies with invertebrate models of disease.

## MATERIALS AND METHODS

### Fly Husbandry and Stocks

Flies were maintained on a cornmeal, molasses, and yeast diet at 21⁰C in wide vials unless otherwise noted. For antibiotic feeding, standard food was melted and mixed with the antibiotics once it cooled to 50-60⁰C. Antiobiotic cocktail #1 contained metronidazole (150µg/mL), ampicillin (50µg/mL), erythromycin (50µg/mL), carbenicillin (150µg/mL), and kanamycin (50µg/mL). Antiobiotic cocktail #2 contained carbenicillin (150µg/mL), metronidazole (150µg/mL), and tetracyclin (75µg/mL). Antibiotic food was stored for a maximum of two weeks prior to use and flies were flipped onto new food each day. A complete list of stocks used is given in the Key Resources table.

### Ovarian Tumor Induction and Transplantation

After eclosion, adult females were raised at 21⁰C on food with yeast powder for two days. No more than 20 flies were kept in each wide vial. Flies were then moved to new food and shifted to 29⁰C to initiate tumor induction. Non-tumor bearing flies were treated identically in parallel to control for the temperature change.

For ovarian tumor transplants, our technique was adapted from a previously published protocol (Rossi & Gonzalez, 2015). Tumors were induced in flies expressing *aPKC^ΔN^, Ras^V12^* and *His2av::RFP* under control of *tj-GAL4, tub-GAL80^ts^* for 8-12 days. This timing was selected to allow donor flies to develop grade 2 tumors (see Hsi et al 2023 for tumor grading criteria). Individual tumorous ovarioles were dissected and carefully transferred into the abdominal cavity of recipient flies with a sharpened glass micropipette <200µm in diameter. After surgery, flies were allowed to recover for at least 30 minutes at room temperature before being placed at 29⁰C. For recipient hosts in which RNAi knockdown is performed, flies were kept at 29⁰C for 3-4 days prior to performing transplantation on them. Successful grafting was determined by the presence of red epifluorescence in the abdomen 6 days after surgery or by identifying tumor growth in the abdomen upon necropsy. Flies with tumors grafting on the gut itself were excluded from experiments.

### Lifespan Assays

Food was changed every two days and the number of dead flies was counted. For each lifespan assay, at least 55 flies were used for each sample group, with control and experimental groups assessed contemporaneously. RNAi stocks were backcrossed at least four generations to an isogenized *w^1118^* line (BDSC #5905) to minimize variation in genetic background.

### Gut Microbe CFU Assay

Up to 25 flies were washed in an Eppendorf with 1mL of 70% ethanol by inverting for at least two seconds. The ethanol wash was performed twice before transferring flies to a clean Kimwipe to dry. 3 clean flies were placed in an Eppendorf with 150µL of sterile PBS and homogenized with a plastic pestle for at least 60 seconds. The homogenate was serial diluted and 30µL of diluent was plated on 100mm LB agar plates. For each experiment, a plate was set up with 30µL of PBS that was mixed with ethanol-cleaned flies without homogenization to confirm successful surface sterilization. Plates were incubated at 29⁰C for 3-4 days before counting colonies.

### Immunofluorescence

Intestines were dissected in PBS and fixed in 4% PFA-PBS for 40 minutes at 21⁰C. To image gut-associated macrophages, the intestine was fixed while kept in the opened abdomen to minimize macrophage detachment. Dorsal fat body, thoracic muscle and ovaries were fixed for 1 hour in 4% PFA-PBS at 21⁰C. After washing three times with PBS, samples were blocked for at least 30 minutes with 4% NGS/1% BSA in PBST (0.2%). Samples were incubated overnight at 4⁰C in primary antibodies diluted in block as follows: Dlg (1:200), Coracle (1:50), cleaved DCP1 (1:200), GFP (1:1000) and dpERK (1:200). For dpERK stains, samples were fixed for 30 minutes and then subjected to a serial methanol dehydration. Fixation, dehydration, and primary antibody incubations were performed with the addition of phosphatase inhibitor (1:100, Sigma P5726). Samples were incubated in Alexafluor-conjugated secondary antibodies in block for 2 hours at 21⁰C or overnight at 4⁰C. To stain F-actin, fixed tissues were incubated for 20 minutes at room temperature with fluorophore-conjugated phalloidin at a concentration of 1:200. To stain nuclei, a 1:1000 DAPI solution was applied for 20 minutes. Following all antibody and stain incubations, samples were washed three times with PBS and left in the final wash for at least 20 minutes. Samples were mounted in 0.5% w/v Propyl Gallate in 1:1 PBS-Glycerol. All samples were imaged on a Zeiss LSM900 Scanning Confocal with a Plan-APOCHROMAT 20x/0,8 objective.

### Ex-Q Assay

We modified the protocol for the Ex-Q assay from a previously published protocol(Wu *et al*, 2019). One 5mL dual-position culture tube was prepared per fly assayed. Using a sterile needle, holes were poked into the plastic ring around the top exterior of the cap to allow air extra air ventilation. The interior of the cap has a small cup. Melted food with 1% erioglaucine dye is pipetted into the cup and allowed to set. Individual flies were placed into culture tubes with caps containing dyed food and kept at 29⁰C for 24 hrs within a humidified container. Then, the cap was replaced with a new cap containing undyed food and the flies were incubated for an additional 3 hours at 29⁰C in order to allow residual dyed food to empty from the digestive tract. The flies were then removed and the cap were again swapped out for caps without food. 500µL of PBS was added to the culture tube and the tube was vortexed and nutated thoroughly to fully dissolve the dye-containing excretions. The absorption at 630nm was then quantified for each sample. To blank the spectrophotometer, PBS added to an empty culture tube that had been incubated along side the experimental samples was used.

### CAFE Assay

We modified the protocol for the CAFE assay from a previously published protocol (Diegelmann *et al*, 2017). A custom cap fitted with a rubber o-ring was made to a fit a 3.8cm diameter plastic vial. The vials were 7cm tall. The cap contained 6 small holes to fit glass capillaries and a central hole to fit a moistened piece of sponge. Two layers of moistened filter paper were placed at the bottom of the vial. Glass capillaries were filled with 50% sucrose solution tinted with a small amount of Red #40 Dye.

Filled capillaries were inserted into the cap such that the end of the capillary floated in the middle of the vial. 7-20 flies were kept in each vial. Prior to transferring flies into prepared vials, animals were pre-starved for 3 hours in empty vials containing moistened filter paper to maintain their hydration. For every experiment, an additional vial was prepared containing no flies as a control for evaporation of the sucrose solution. Vials were transferred to a large Tupperware container kept humid with small vessels of water. This container was incubated for 24 hours at 29⁰C. The sucrose solution depleted from each capillary was measured and the total loss of fluid from evaporation was deducted from these values. Then, the amount of food eaten per fly was calculated for each vial.

### Smurf Assay

The smurf assay was performed as described previously (Martins *et al*, 2018). Briefly, Blue Dye #1 was mixed with melted molasses food to a final concentration of 2.5% w/v. 6 days after shifting to 29⁰C, flies were transferred to vials of blue food. A maximum of 8 flies were housed per vial. Every other day, flies were transferred to fresh blue food. 6 days per week, flies were scored for blue color outside of the gut tract and death.

### Image Analysis

To measure the Dlg plasma membrane index, a maximum project of 1-3 slices was made at the level of the septate junctions of the epithelium. A 3-pixel wide segmented line traced along an interface of cell junctions oriented radially with the lumen. The mean gray value of the junctions (P) was measured. The segmented line ROI was shifted 5-10 pixels away such that it overlayed the cytoplasm of the cells and a second mean gray value was obtained (C). The plasma membrane index is calculated as P ÷ C. This measurement was repeated across three interfaces evenly spaced across the R5 region and these values were averaged to give a final measurement for the sample. For fluorescent signal intensity, mean gray values were obtained from maximum projected images after selecting the region of interest with the polygon selection tool in FIJI. In the ovary, measurements were taken across approximately the anterior half of the ovary. For the fat body, the entire dissected dorsal cuticle was measured excluding regions of cuticle overlap. *NRE+* and *esg*+ cells were counted in the R5 gut region in the half of the epithelium closest to the microscope objective. The spot detection function in Imaris 10.2 software was used to count cells expressing GFP markers in the R5 region of the intestine.

## Supporting information

Supplemental Info Combined

## ACKNOWLEDGEMENTS

We thank Brian Stramer, Denise Montell and Lucy O’Brien for generous gifts of reagents, as well as the community resources provided by the Bloomington Drosophila Stock Center (NIH P40OD018537), TRiP at Harvard Medical School (NIH/NIGMS R01-GM084947), the Vienna Drosophila Resource Center and the Kyoto stock center. We acknowledge Lucy O’Brien and her lab members for feedback and guidance on this work. We are grateful to the entire Bilder lab for helpful discussions. Imaris software access and training was provided by the UC Berkeley Cancer Research Laboratory’s Molecular Imaging Center. This work was supported by a Mark Foundation ASPIRE Award, American Cancer Society grant DBG-24-1322649, and NIH grant GM130388 to D.B., as well as a Damon Runyon – Mark Foundation Postdoctoral Fellowship 2400-20 to K.O.

## AUTHOR CONTRIBUTIONS

K.O. and D.B. designed the study and wrote the manuscript. K.O performed most of the experiments, with the help of K.A., and created the figures. J.C. developed and performed the ovarian transplant protocol.

