## Supplemental Info Combined for "A chronic interorgan wound response appropriated by *Drosophila* tumors to induce intestinal inflammation"

**Supplementary Figure 1. Ovarian tumor graft remotely induces intestinal dysplasia.**

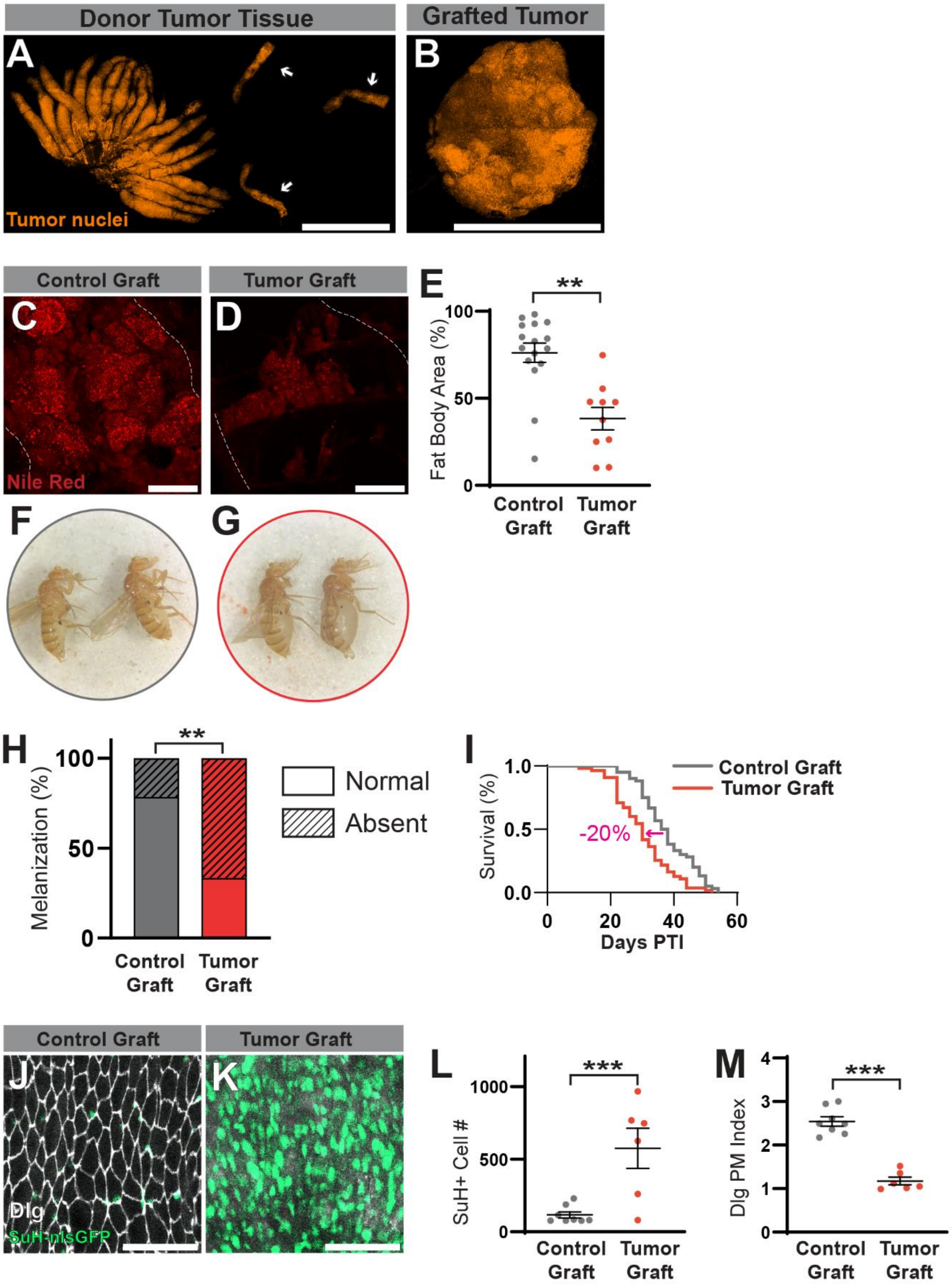


**Supplementary Figure 1: Ovarian tumor graft remotely induces intestinal dysplasia** (**A**) Dissected tumorous ovary with individual ovarioles (white arrows) prepared for transplantation procedure. Tumor cells express His2av::RFP (orange). (**B**) Successfully grafted ovarian tumor dissected from a host fly, 19 days after transplant. (**C**-**E**) Flies with ovarian tumor grafts experience fat body wasting, quantified in (**E**). (**F**, **G**) Tumor grafting induces fluid retention and bloating. (**H**) Tumor-bearing flies lose the capacity for wound melanization. (**I**) Lifespan is reduced with tumor grafting. (**J**-**M**) Dysplasia characterized by SJ disorganization and EB expansion is observed in tumor-bearing flies. Quantified in (**L**, **M**). Scale bars in **A**, **B** = 1000μM. Scale bars in **C**, **D** = 200μM. Scale bars in **J**, **K** = 50μM. Error bars = S.E.M. Log-rank test was used to assess statistical significance in (**K**). Student's t-test was used to assess statistical significance in all other graphs.

**Supplementary Figure 2: Dysplasia caused by tumors is independent of cachexia, feeding behavior and microbiome load.**

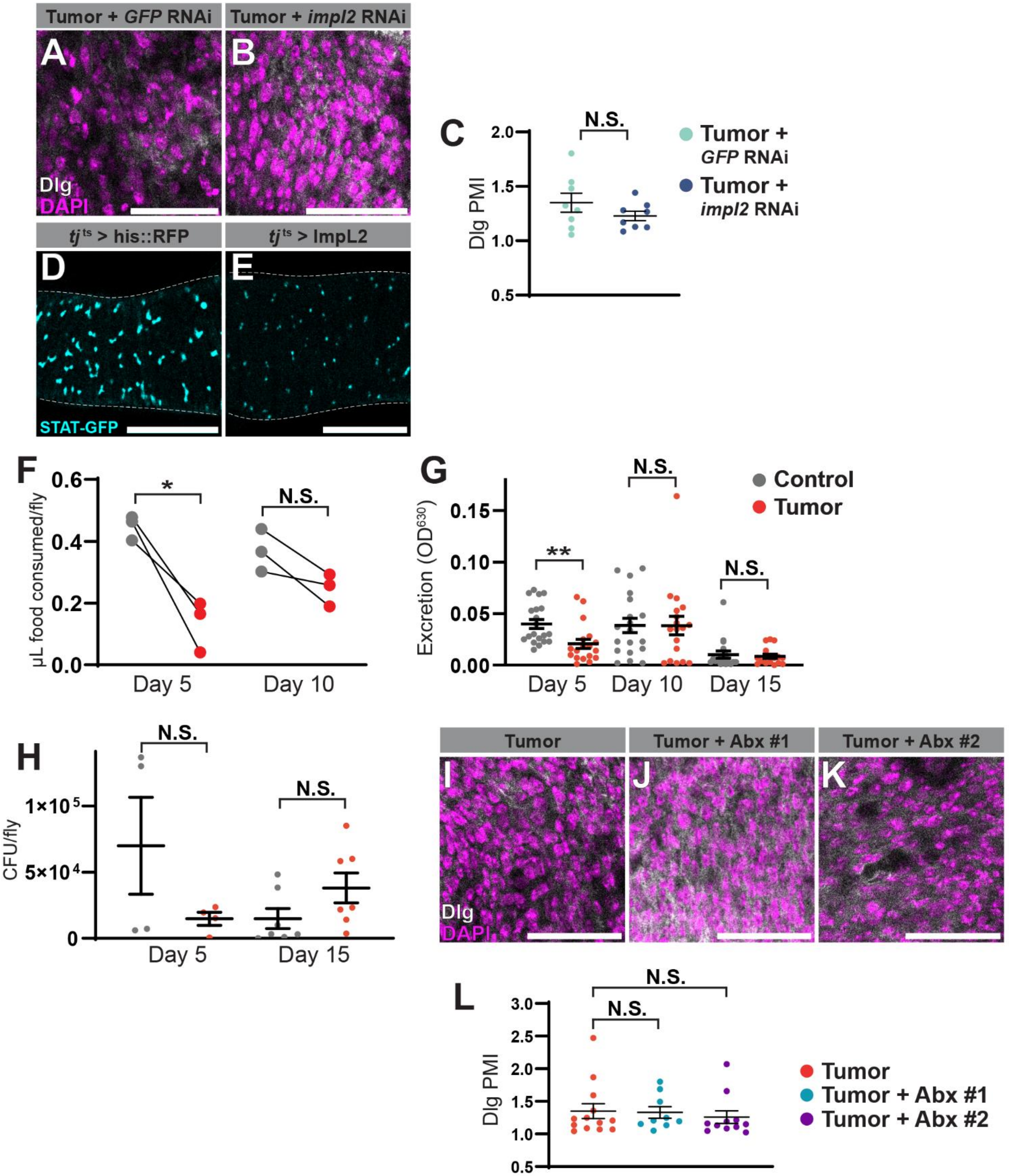


**Supplementary Figure 2: Dysplasia caused by tumors is independent of cachexia, feeding behavior and microbiome load** (**A**-**C**) Knockdown of tumor *ImpL2* does not rescue SJ disorganization or cellular density at 20 days ATI. SJ integrity measured in (**C**). (**D**, **E**) Systemic wasting caused by ImpL2 overexpression does not activate ISC proliferation, as observed by a STAT activity reporter (cyan) to mark ISCs and progenitors. (**F**, **G**) Food consumption as measured by CAFE assay (**F**) and Ex-Q assay (**G**) was only slightly lower in tumor-bearing flies at 5 days ATI. (**H**) Microbial load is not significantly different between control and tumor-bearing flies. (**I**-**L**) Elimination of the gut microbiome did not restore SJ integrity or cellular density. Two different antibiotic combinations (#1 and #2) were used to eliminate the gut microbes (see Materials and Methods). Quantification in (**L**). Scale bars **A**, **B**, **I**-**K** = 50μm. Scale bars **D**, **E** = 100μm. Error bars = S.E.M. Student's t-test used to assess statistical significance in all graphs.

**Supplementary Figure 3. Ovarian tumors and chronic injuries remotely induce inflammatory changes in the intestine.**

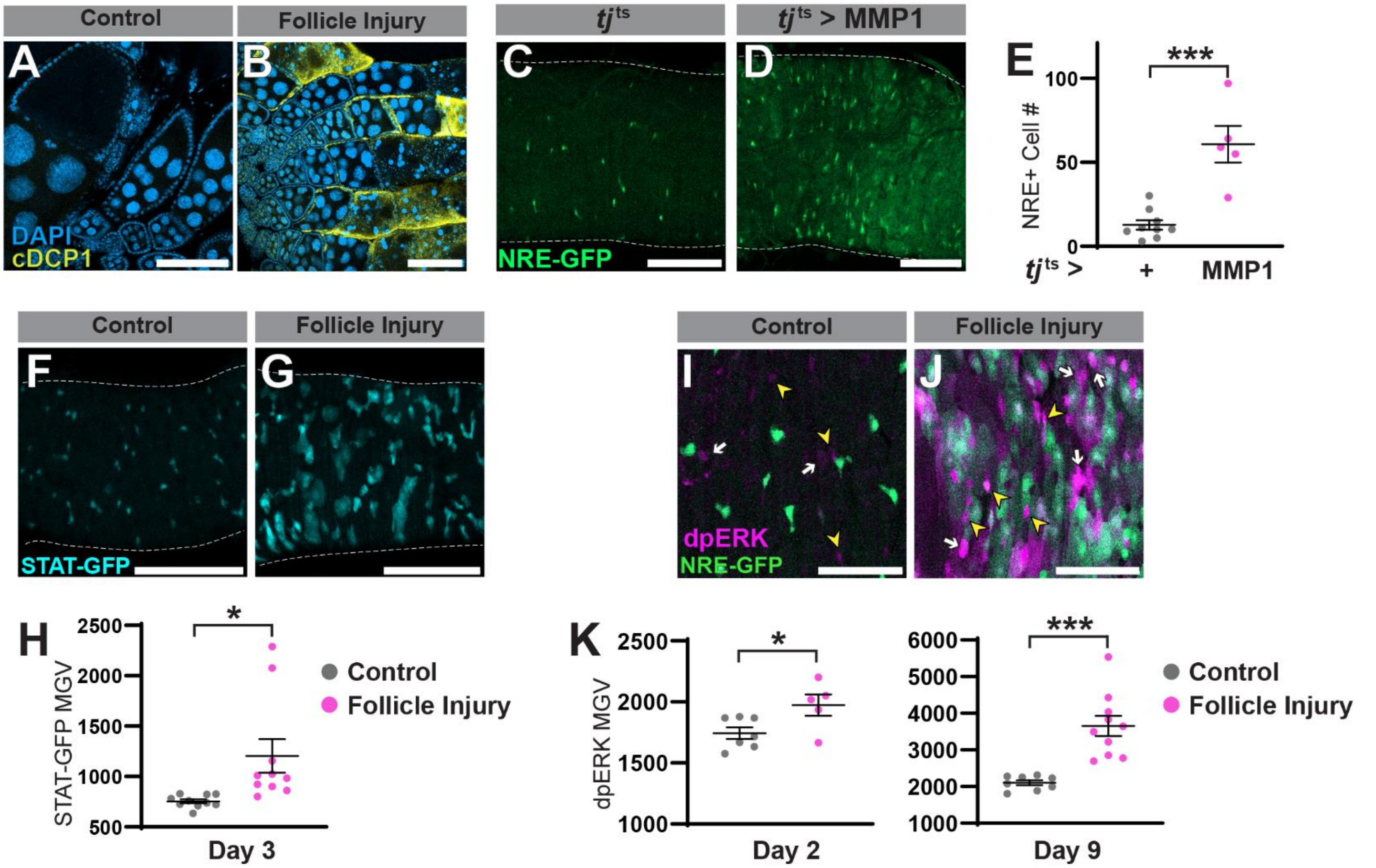


**Supplementary Figure 3. Ovarian tumors and chronic injuries remotely induce inflammatory changes in the intestine.** (**A**, **B**) *rpr* expression in the ovarian epithelium leads to widespread cell death as determined by cleaved DCP-1 staining (cDCP1, yellow). (**C-E**) MMP1 overexpression in the ovarian epithelium results in increased NRE-GFP+ EBs (green). Quantified in (**E**). (**F-H**) STAT-GFP reporter (cyan) shows activation in the gut epithelium upon chronic ovarian injury, quantified in (**H**). (**I-K**) dpERK (magenta) staining revealed MAP Kinase activation in the intestine following chronic ovarian injury, quantified in (**K**). Scale bars in **A-D**, **F**, **G** = 100µm. **J**, **K** = 50µm. Error bars = S.E.M. Student's t-test used to assess statistical significance in all graphs.

**Supplementary Figure 4. Tumor-driven coagulopathy engages tissue-specific, chronic inflammation**.

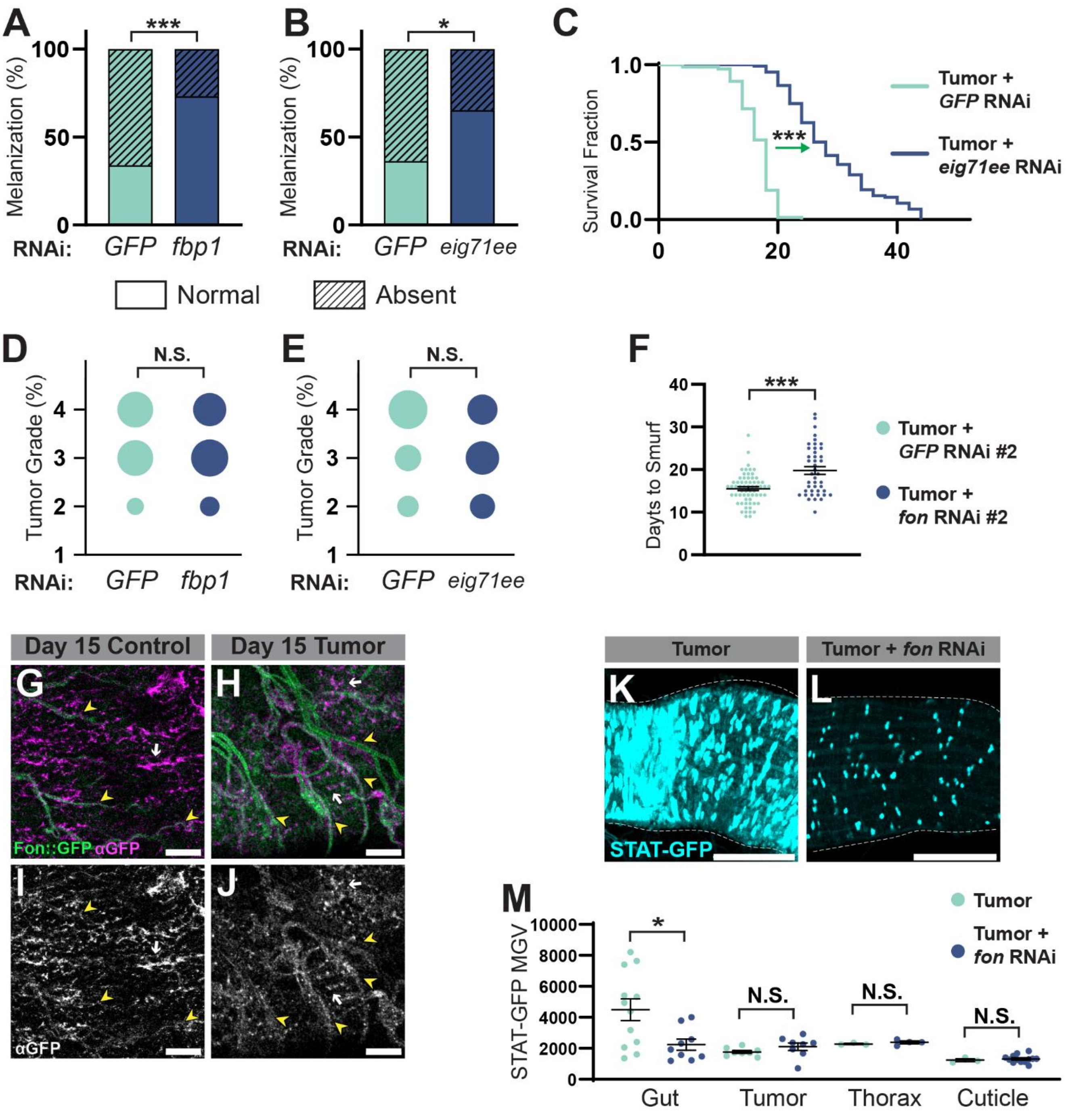


**Supplementary Figure 4. Tumor-driven coagulopathy engages tissue-specific, chronic inflammation**. (**A**-**B**) Clotting components Fbp1 and Eig71ee regulate loss of melanization in tumor-bearing flies. (**C**) Knockdown of *eig71ee* in the tumor extends lifespan. (**D**, **E**) Tumor grade is unaffected by *fbp1* nor *eig71ee* knockdown. (**F**) A second RNAi against *fon* delays smurfing onset when expressed in the tumor. (**G**-**H**) Fon::GFP is recruited to gut muscle (white arrow) and trachea. (yellow arrowheads). Intrinsic GFP signal is displayed in green while extracellular anti-GFP immunofluorescence is shown in magenta. Trachea signals are qualitatively more prominent in tumor-bearing flies. (**K**-**L**) Fon knockdown selectively attenuates JAK/Stat activation (cyan) in the intestine. Quantification of intestine and other tissues in (**M**). Scale bars in **G**-**J** = 25μm. Scale bars in **K**, **L** = 100μm. Error bars = S.E.M. Log-rank test was used to assess statistical significance in (**C**). Student's t-test used to assess statistical significance in all other graphs.

### Key resources table

| REAGENT or RESOURCE | SOURCE | IDENTIFIER |
|---|---|---|
| **Antibodies** | | |
| Anti-Discs large (mouse) | Developmental Studies Hybridoma Bank | 4F3 |
| Anti-cleaved DCP1 (rabbit) | Cell Signaling Technologies | 9578S |
| Anti-Coracle (mouse) | Developmental Studies Hybridoma Bank | C615.16 |
| Anti-GFP (chicken) | Invitrogen | A10262 |
| Anti-diphosphoERK (rabbit) | Cell Signaling Technologies | 4370S |
| **Biological samples** | | |
| none | | |
| **Chemicals, peptides, and recombinant proteins** | | |
| TRITC-Phalloidin | Sigma-Aldrich | P1951 |
| DAPI | ThermoFisher Scientific | D1306 |
| Nile Red | Sigma-Aldrich | N3013 |
| Paraformaldehyde 16% | Electron Microscopy Sciences | 15710 |
| Triton-X | Sigma-Aldrich | T8787 |
| Phosphate Buffered Solution | Sigma-Aldrich | P4417 |
| Bovine Serum Albumin | Sigma-Aldrich | A3912 |
| Alexa Fluor Phalloidin 647 | Invitrogen | A22287 |
| Alexa Flour Phalloidin 488 | Invitrogen | A12379 |
| Normal Goat Serum | Celltech | 5425 |
| Ampicillin | Sigma Aldrich | A91518 |
| Carbenicillin | bioWORLD | 40310030-1 |
| Tetracycline Hydrochloride | Sigma-Aldrich | T7660 |
| Metronidazole | Research Products International | M81000 |
| Kanamycin Sulfate | Gibco | 15160-054 |
| Erythromycin | Sigma-Aldrich | E5389 |
| Red Dye #40 | Spectrum Chemical | FD140 |
| Erioglaucine Sodium Salt | Thomas Scientific | J61780 |
| Phosphatase Inhibitor Cocktail 2 | Sigma-Aldrich | P5726 |
| **Critical commercial assays** | | |
| none | | |
| **Deposited data** | | |
| none | | |
| **Experimental models: Cell lines** | | |
| none | | |
| **Experimental models: Organisms/strains** | | |
| *Drosophila: tj-GAL4* | Kyoto Stock Center | 104055 |
| *UAS-aPKC*$^{\Delta N}$ | Bloomington Drosophila Stock Center | 51673 |

| | | |
|---|---|---|
| *UAS-Ras$^{V12}$* | Bloomington Drosophila Stock Center | 4847 |
| *Tub-GAL80$^{ts}$* | Bloomington Drosophila Stock Center | 7109 |
| *UAS-Fon::GFP* | Bloomington Drosophila Stock Center | 43646 |
| *P{VDRCsh60200}attP40* | Vienna Drosophila Resource Center | 60200 |
| *UAS-GFP #1 RNAi* | Vienna Drosophila Resource Center | 60201 |
| *UAS-Fon #1 RNAi* | Vienna Drosophila Resource Center | 330575 |
| *UAS-Fon #2 RNAi* | Hsi *et al*, 2023 | FBal0400664 |
| *UAS-FBP1 RNAi* | Vienna Drosophila Resource Center | 330200 |
| *UAS-ImpL2 RNAi* | Bloomington Drosophila Stock Center | 64936 |
| *UAS-Eig71Ee RNAi* | Vienna Drosophila Resource Center | 330464 |
| *UAS-EGFR RNAi* | Bloomington Drosophila Stock Center | 25781 |
| *UAS-Stat92E RNAi* | Bloomington Drosophila Stock Center | 33637 |
| *UAS-Dome RNAi* | Bloomington Drosophila Stock Center | 34618 |
| *UAS-Stg RNAi* | Bloomington Drosophila Stock Center | 34831 |
| *esg-GAL4* | Kyoto Drosophila Stock Center | 112304 |
| *GBE-su(H)nls::GFP* | de Navascués *et al*, 2012 | PMID: 22522699 |
| *UAS-his2b::CFP* | Yoshihiro Inoue Lab | N/A |
| *NRE-GFP* | Bloomington Drosophila Stock Center | 30728 |
| *esg-sfGFP* | Bloomington Drosophila Stock Center | 78333 |
| *LexAop-Ras$^{V12}$* | Adiga *et al*, 2025 | N/A |
| *LexAop-aPKC$^{\Delta N}$* | Adiga *et al*, 2025 | N/A |
| *tj-LexA::GAD* | Adiga *et al*, 2025 | N/A |
| *IDGF3::GFP* | Kucerova *et al*, 2015 | FBal0327371 |
| *Mdu::GFP* | Bloomington Drosophila Stock Center | 50849 |
| *LexAop-Rpr* | Santabárbara-Ruiz *et al*, 2015 | FBal0346745 |
| **Oligonucleotides** | | |

| | | |
|---|---|---|
| none | | |
| **Recombinant DNA** | | |
| none | | |
| **Software and algorithms** | | |
| Imaris 10.2 | Oxford Instruments | imaris.oxinst.com |
| FIJI | ImageJ | fiji.sc |
| Prism v10 | GraphPad | graphpad.com |
| Zeiss Imaging Software (Zen) | Zeiss | zeiss.com/microscopy/us/products/microscope-software/zen.html |
| **Other** | | |
| Blaubrand 5μL micropipettes with ring marks | Brand | 708707 |
